# The N-coil and the globular N-terminal domain of plant ARGONAUTE1 are interaction hubs for regulatory factors

**DOI:** 10.1101/2023.01.18.524620

**Authors:** Simon Bressendorff, Swathi Kausika, Ida Marie Zobbe Sjøgaard, Emilie Duus Oksbjerg, Alec Michels, Christian Poulsen, Peter Brodersen

**Affiliations:** University of Copenhagen, Copenhagen Plant Science Center, Ole Maaløes Vej 5, DK-2200 Copenhagen N, Denmark; Novo Nordisk A/S, Novo Nordisk Park 1, DK-2760 Måløv, Denmark

**Keywords:** ARGONAUTE, N domain, N-coil, regulated proteolysis, F-box protein, autophagy cargo receptor, ATI1, ATI2

## Abstract

The effector complex of RNA interference (RNAi) contains at its core an ARGONAUTE (AGO) protein bound to a small guide RNA. AGO proteins adopt a two-lobed structure in which the N-terminal (N) and Piwi-Argonaute-Zwille (PAZ) domains make up one lobe, while the middle (MID) and Piwi domains make up the other. Specific biochemical functions of PAZ, MID and Piwi domains of eukaryotic AGO proteins have been described, but the functions of the N-terminal domain remain less clear. Here, we use yeast two-hybrid screening with the N-terminal domain of the founding member of the AGO protein family, arabidopsis AGO1, to reveal that it interacts with many factors involved in regulated proteolysis. Interaction with a large group of proteins, including the autophagy cargo receptors ATI1 and ATI2, requires residues in a short, linear region, the N-coil, that joins the MID-Piwi lobe in the three-dimensional structure of AGO. In contrast, the F-box protein AUF1 interacts with AGO1 independently of the N-coil and requires distinct residues in the globular N domain itself. Mutation of AGO1 residues necessary for interaction with protein degradation factors in yeast stabilizes reporters fused to the AGO1 N-terminal domain in plants, supporting their *in vivo* relevance. Our results define distinct regions of the N domain implicated in protein-protein interaction, and point to a particular importance of the AGO1 N-coil as a site of interaction with regulatory factors.

## INTRODUCTION

ARGONAUTE (AGO) proteins are the central protein components of RNA Induced Silencing Complexes (RISCs) whose activities determine all RNA interference (RNAi) phenomena. RISCs use base pairing of a small non-coding RNA bound to AGO as a specificity determinant in the selection of complementary target RNA for regulation. This principle of RNA regulation underlies fundamental elements of endogenous gene regulation and defense against parasitic genetic material such as transposable elements and viruses.

In higher plants, many distinct AGO paralogues perform functions in different RNAi pathways. For example, the founding member of the AGO protein family, arabidopsis AGO1, carries out post-transcriptional gene regulation. This includes both endogenous gene regulation via the majority of microRNAs (miRNAs) and some small interfering RNAs (siRNAs), and antiviral defense via virus-derived siRNAs [1, 2]. Other AGO proteins, including AGO2, AGO7 and AGO10, have additional functions in antiviral RNAi [3–7] and endogenous gene regulation, sometimes in a highly specialized manner that relies on preferential association with one or only few miRNAs [1, 8–10]. A separate phylogenetic clade of AGO proteins, represented by AGO4, AGO6 and AGO9 in arabidopsis, acts as effectors of RNA-directed DNA methylation that silences transposable and other repetitive DNA elements at the transcriptional level [11].

Despite their different biological functions, AGO proteins have a recurrent domain architecture and a highly conserved two-lobed three-dimensional structure when bound to small RNA [12–15]. Four domains are recurrent: The N-terminal (N) domain, the PAZ (Piwi-Argonaute-Zwille) domain, the MID (middle) domain, and the Piwi domain [16]. The N and PAZ domains form one lobe and are connected by a linker region referred to as L1, while MID and Piwi form the other lobe. The PAZ and MID domains are connected by a long linker region called L2 that interacts extensively with both the small guide RNA and with other domains in AGO (Figure 1). The MID domain harbors a pocket dedicated to the tight binding of both the 5’-phosphate and the nucleobase of the 5’-nucleotide of the small guide RNA [17, 18]. In contrast, the PAZ domain binds the 3’-end [19] in such a way that it can be released to participate in base pairing to a target RNA [20, 21], a process that requires extensive conformational rearrangement of the AGO-small RNA complex [22, 23]. The Piwi domain adopts a ribonuclease H-like fold [24], and, accordingly, contains the two-metal ion catalytic center of those AGO proteins capable of cleaving target RNAs [14]. In addition, the Piwi domain of many AGO proteins has hydrophobic binding pockets that accommodate indole side chains of AGO interactors that may be crucial for RISC activity [13, 25].

**Figure 1.**
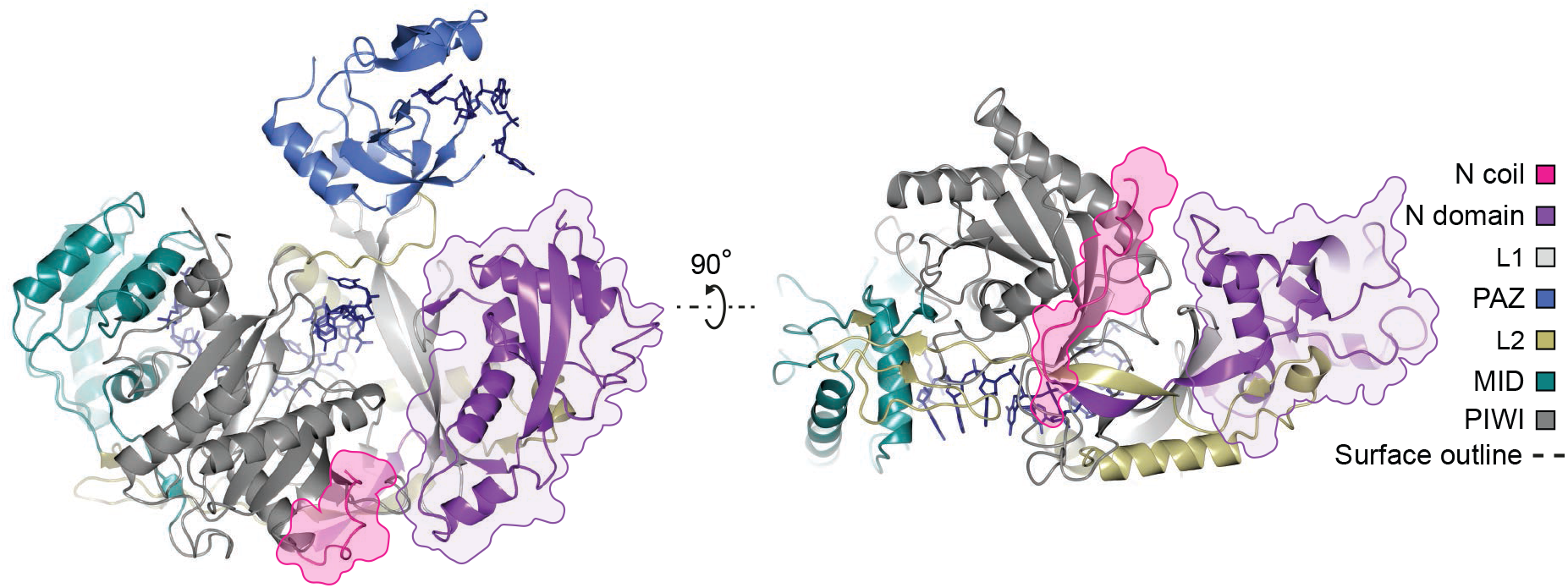
Overview of AGO protein structure. Structure of human Ago2 with its different domains and linker regions shown in distinct colors as indicated in the caption on the right. The N-coil associated with the MID-Piwi lobe is highlighted in pink, and globular N-domain in the N-PAZ lobe is highlighted in purple. The presentations were made using the coordinates from the crystal structure of the miR20a-bound human Ago2 (PDB 4F3T)[15].

In contrast to the PAZ, MID and Piwi domains, the biochemical functions of the N domain remain less well defined. Studies of RISC loading using the fly Ago1 protein suggested a role of the N domain in small RNA duplex separation [26], but the mutations with the strongest defect in duplex separation mapped to the L1 region rather than the N domain itself. This is consistent with the phenotype of arabidopsis mutants homozygous for the *ago1-57* allele that encodes a point mutant in L1 (G371D) with a specific defect in unwinding of duplex siRNAs [27]. Studies of human Ago proteins identified a region in the N-terminal part of Ago2 that was sufficient to confer catalytic activity to Ago3 upon replacement of the corresponding part of Ago3 [28], perhaps suggesting an implication of this part in guiding the structural rearrangements required to reach the catalytically active conformation [22]. Interestingly, although this part of human Ago2 is located towards the N-terminus in the primary structure, it is not fully contained in the N-PAZ lobe in the threedimensional structure of the Ago2-small RNA complex (Figure 1). A 14-amino acid linear segment interacts with the Piwi domain and is part of the MID-Piwi lobe, while two β-strands connects it to the globular N domain. We previously described the 14-amino acid segment as a structural unit of AGO proteins, and termed it the N-coil [1]. The N-coil also stands out in collections of arabidopsis *ago1* mutant alleles defective in miRNA function, because it is the only part of the N-terminal region in which point mutations are known to cause reduction of miRNA function [1]. Thus, there is molecular genetic evidence that the N-coil is important for AGO function, but few hints from genetic or molecular studies as to what functions the globular N-domain may have.

In this study, we identify interactors of the N-coil-Globular N (NcGN) part of arabidopsis AGO1 using yeast two-hybrid screening. We show that factors implicated in regulated proteolysis are prominent among such interactors, and that two types of interactors can be defined. The first and largest group contains interactors whose binding to the NcGN requires an intact N-coil. The second type interacts with the NcGN independently of the N-coil, but requires specific residues in the GN for binding. These results define at least two distinct modes by which the N-terminal part of AGO1 interacts with regulatory factors, in particular factors involved in regulated proteolysis.

## RESULTS

### The N-coil-Globular N part of AGO1 interacts with multiple factors implicated in regulated proteolysis

To reveal functions of the poorly understood N-coil-Globular N part of arabidopsis AGO1 (NcGN^AGO1^), we first conducted yeast two-hybrid screens with NcGN^AGO1^ as a bait against three prey cDNA libraries. This effort yielded 15 potential NcGN^AGO1^ interactors, validated in the two-hybrid system by isolation of the prey plasmid and re-transformation with the NcGN^AGO1^ bait into the reporter strain (Table 1, Supplementary Figure S1). Several candidate interactors had predicted biochemical functions related to regulated proteolysis via either the ubiquitin-proteasome pathway or via autophagy (Table 1). This also included the two intrinsically disordered, transmembrane autophagy cargo receptor proteins ATI1 and ATI2 [29, 30] that were previously shown to be implicated in degradation of AGO1 via the poleroviral RNAi suppressor P0 [31]. The trend towards protein degradation was not exclusive, however, as other interactors were also recovered, including Hsp90, a protein phosphatase, and vesicle trafficking and membrane contact proteins (VAP27, NTMC2T5.1) [32, 33] (Table 1).

**Table 1.** List of genes exhibiting positive yeast two-hybrid interaction with NcGN^AGO1^. The two-hybrid interactions were confirmed by retransformation of the plasmid recovered in the large-scale screen with AGO1 NcGN bait plasmid. CD4-22, yeast two-hybrid cDNA library prepared from RNA from 3-day old etiolated Arabidopsis (Col-0) seedlings [51]; CD4-10; yeast two-hybrid cDNA library prepared from RNA from Arabidopsis, ecotype Nössen; MM, Matchmaker Arabidopsis yeast two-hybrid cDNA library produced and sold by Clontech ®. ATI1 was not recovered in any of the large-scale screens, but tested positive in a directed two-hybrid assay designed because of its similarity to ATI2. IDR, intrinsically disordered region; TM, transmembrane domain; DUF, domain of unknown function; CC, coiled-coil; ZnF, Zinc finger; UBL, ubiquitin-like; ATPase, ATP hydrolyzing domain of Hsp90; TCTP, Translationally Controlled Tumor Protein; EF-hand, Ca^2+^-binding EF-hand; MSP, nematode Major Sperm Protein immunoglobulin-like domain; C2, Ca^2+^-dependent membrane-targeting domain; PP2C, Metal ion-dependent protein phosphatase; SOD, superoxide dismutase.

|  | Library | AGI | Gene name | Function | Size | Interacting region | Domains in interacting piece |
| --- | --- | --- | --- | --- | --- | --- | --- |
| 1 | CD4-22 | AT4G00355 | ATI2 | ATG8-interacting | 266 | 1-267 | IDR, TM |
| 2 | Directed | AT2G45980 | ATI1 | ATG8-interacting | 256 | 1-256 | IDR, TM |
| 3 | CD4-22 | AT1G78100 | AUF1 | F-box | 334 | 85-334 | DUF |
| 4 | CD4-22 | AT2G32560 | AUF3 | F-box | 371 | 171-371 | DUF |
| 5 | CD4-22 | AT2G44950 | HUB1 | RING finger | 878 | 721-878 | CC, ZnF |
| 6 | CD4-22 | AT5G25270 | Ub-like | Ubiquitin-like | 642 | 1-253 | UBL, IDR |
| 7 | CD4-22 | AT5G56010 | HSP90.3 | Heat Shock Protein 90 | 699 | 81-208 | ATPase |
| 8 | CD4-22 | AT3G16640 | TCTP | Translationally controlled tumor protein | 168 | 1-168 | TCTP |
| 9 | CD4-22 | AT5G21274 | CAM6 | Calmodulin | 149 | 1-149 | EF-hand |
| 10 | CD4-22 | AT3G60600 | VAP27 | VAMP-associated protein; vesicle trafficking | 257 | 60-257 | MSP, TM |
| 11 | CD4-22 | AT1G50260 | NTMC2T5.1 | Membrane Contact Site protein | 675 | 414-635 | C2 |
| 12 | MM | AT1G19860 |  | C3H Zn finger | 415 | 144-415 | IDR, ZnF |
| 13 | CD4-22 | AT3G02750 | PP2C | Protein phosphatase | 527 | 47-346 | PP2C |
| 14 | CD4-22/<br>CD4-10 | AT1G07890 | APX1 | Ascorbate peroxidase | 250 | 1-250 | Peroxidase |
| 15 | CD4-22 | AT4G25100 | FeSOD1 | Fe-Superoxide dismutase | 212 | 1-212 | SOD |

### The NcGN of AGO1 functions as a degron

Since “Regulated proteolysis” was the clearest common functional category among candidate interactors, we carried out pulse labeling experiments with yellow fluorescent protein (YFP) and NcGN^AGO1^-YFP fusions stably expressed in *Arabidopsis thaliana* to see whether presence of NcGN^AGO1^ in the YFP fusion protein causes rapid degradation. When lines with comparable YFP steady state levels were analyzed, the synthesis rate (and hence also the degradation rate) of the NcGN^AGO1^-YFP fusion protein was indeed 6.6-fold higher than YFP alone, demonstrating that the NcGN^AGO1^ is subject to rapid proteolysis (Figure 2A,B). We note that recognition of NcGN^AGO1^ by the chaperone-assisted protein degradation machinery as a consequence of its detachment from normally interacting domains in native AGO1 is unlikely, because the Gal4^BD^-NcGN^AGO1^ fusion protein was easily detectable in yeast, and was sufficiently stable for yeast two-hybrid analyses to be carried out (Table 1, Figure 3). Thus, we continued our characterization of NcGN^AGO1^ interactors with an eye towards regulated proteolysis of the AGO1 protein.

**Figure 2.**
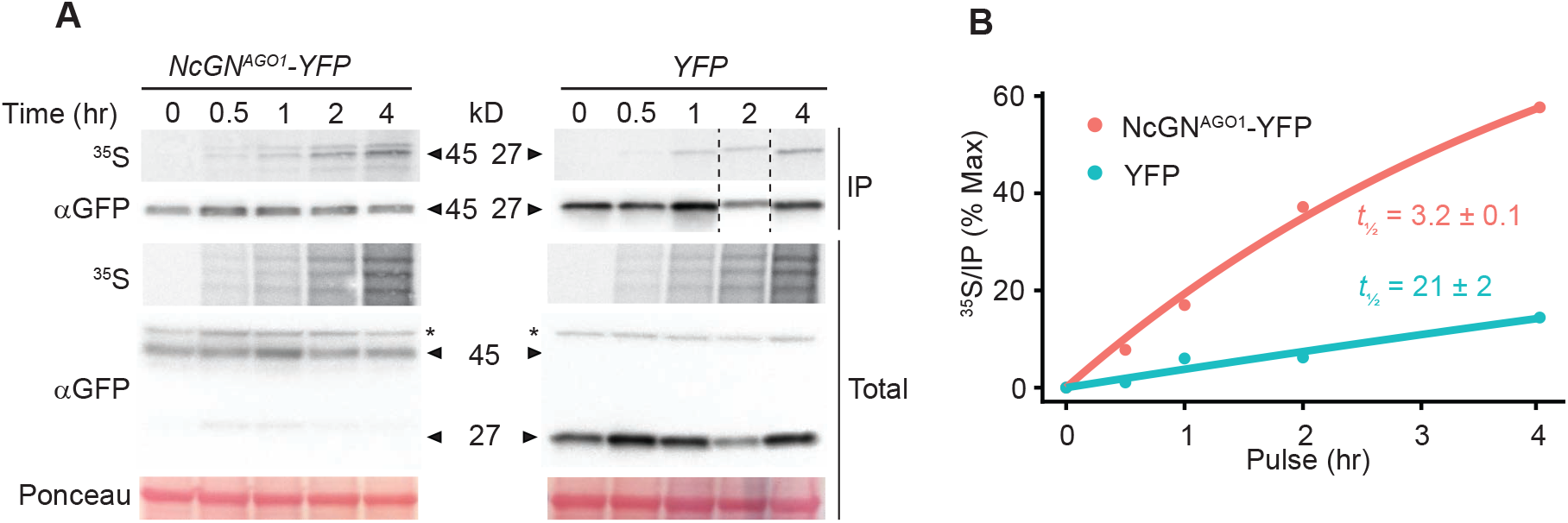
NcGN^AGO1^ is rapidly turned over in stably expressing *Arabidopsis* lines. (A) ^35^S-Met/Cys incorporation into AGO1-NcGN-YFP and YFP, respectively, constitutively expressed in Col-0 WT. 9-day old seedlings were analyzed. Immunoblots and autoradiograms are shown both for analysis of α-GFP immuno-affinity purified fractions and for total lysates. The dashed lines indicate lane cropping to correct the loading order for presentation purposes. Ponceau staining shows total protein. The * indicates unspecific bands. (B) Quantification of ^35^S-Met/Cys incorporation into α-GFP immunoprecipitated protein as a function of time. Data are fitted to an exponential model reporting *t_½_* in hours +/- standard deviation, as detailed in Methods.

**Figure 3.**
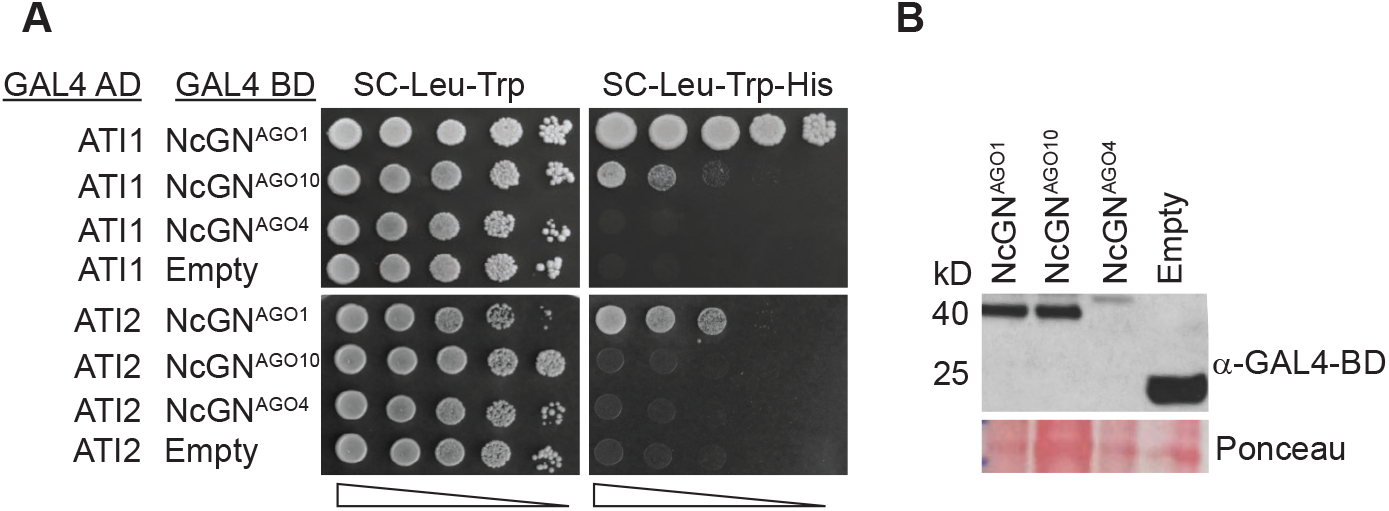
ATI1 and ATI2 interact specifically with NcGN^AGO1^. (A) Yeast two-hybrid analysis of ATI1 and ATI2 fused to the activation domain (AD) of the yeast transcription factor Gal4 co-expressed either with NcGN domains of AGO1, AGO10, or AGO4 fused to the DNA binding domain of Gal4 (BD), or with Gal4-BD alone (empty). 10-fold serial dilutions of yeast cells were spotted on permissive (left) and selective (right) media. (B) Immunoblot analysis with antibodies recognizing the DNA binding domain (α-GAL4-BD) of yeast strains expressing ATI2 fused to the activation domain (AD) of Gal4 and NcGN^AGO1^, NcGN^AGO10^ or NcGN^AGO4^ fused to Gal4-BD. Empty vectors expressing unfused Gal4-AD and Gal4-BD were used as negative control. Ponceau shows total protein.

### Most candidate NcGN^AGO1^ interactors are specific for AGO1

The AGO protein family contains 10 paralogs in arabidopsis, of which AGO4 is distantly related while AGO10 is closely related to AGO1. We therefore used the NcGNs of AGO4 and AGO10 to test whether a subset of the identified candidate interactors (ATI1, ATI2, AUF1, AUF3, UBI, HUB1, PP2C and Hsp90.3, see Table 1) were specific to NcGN^AGO1^ or whether they represented generic NcGN^AGO^ binding proteins. Seven of the candidates (ATI1, ATI2, AUF1, AUF3, UBI, HUB1, PP2C) showed yeast two-hybrid interactions preferentially or exclusively with NcGN^AGO1^, while one (Hsp90.3) interacted with the NcGN of all three proteins (Figure 3A,B, Supplementary Figure S2). These results indicate that most of the candidate interactors isolated by yeast two-hybrid screening are specific to AGO1. We also note that the more general NcGN^AGO^ interaction observed with Hsp90.3 (Supplementary Figure S2) is consistent with the involvement of Hsp90 in the loading of different AGO proteins across organisms [34–36].

### Validation of selected interactors with bimolecular fluorescence complementation

To strengthen the evidence that the proteins identified by yeast two-hybrid screening represent *bona fide* AGO1 interactors, we selected a subset (ATI1, ATI2, AUF3, Hsp90.2) of them for interaction tests *in planta* using bimolecular fluorescence complementation (BiFC) with the full-length AGO1 protein. In this set, ATI1 serves as positive control as it has previously been shown to give a positive BiFC interaction with AGO1 [31]. We first conducted a small deletion analysis of the N-terminal intrinsically disordered region (IDR) of ATI2 [30] to identify regions of importance for NcGN^AGO1^ interaction in yeast (Figure 4A). This effort identified a 70 amino-acid region (N118-T193) in the C-terminal part of the IDR as required for NcGN^AGO1^ interaction (Figure 4B). Apart from adding information on the nature of the ATI2-NcGN^AGO1^ interaction, this result allows the design of a suitable negative control for the BiFC experiments. We therefore tested whether the C-terminal half of YFP fused to AGO1 (cYFP-AGO1) would lead to YFP fluorescence emission when coexpressed with the N-terminal half of YFP fused to either ATI1, ATI2, ATI2^Δ119-193^, AUF3^171-371^ or Hsp90. For AUF3, a construct encoding a deletion of the F-box (AUF3^171-371^) was used to avoid rapid degradation of interacting protein *in vivo*, and for Hsp90, the Hsp90.2 (AT5G56030) isoform, nearly identical to Hsp90.3 (AT5G56010) identified in the two-hybrid screen, was used. In all cases, we used the 2in1 BiFC system that expresses both mRFP and cYFP/nYFP fusions from the same plasmid, allowing easy visualization of transient transformation efficiency [37]. These assays showed that YFP fluorescence was specifically detected upon co-expression of cYFP-AGO1 with nYFP-ATI1, nYFP-ATI2, nYFP-AUF3^171-371^, and nYFP-Hsp90.2, but not with the nYFP-ATI2^Δ119-193^ negative control (Figure 5). In addition, compared to AGO1, fluorescence obtained with cYFP-AGO10 and nYFP-ATI1/2 was much weaker, but not abolished (Figure 5), consistent with the residual two-hybrid interaction detected with NcGN^AGO10^ (Figure 3A). These results provide independent evidence that several of the NcGN^AGO1^ interactors identified by yeast two-hybrid screening are genuine AGO1 interactors capable of interacting with the full-length protein when expressed in plant cells.

**Figure 4.**
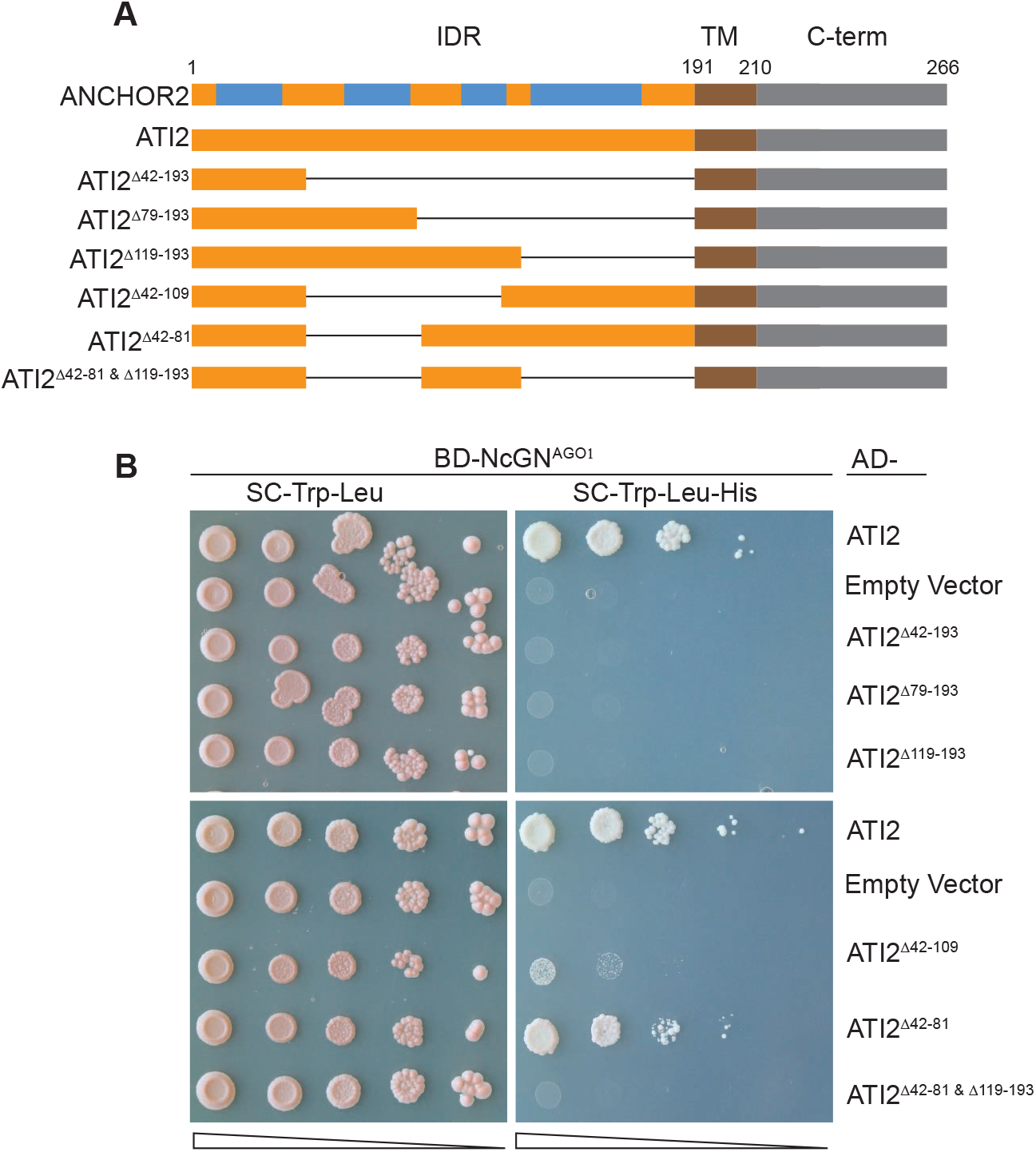
Deletion analysis of the ATI2-NcGN^AGO1^ interaction. (A) Schematic overview of the different ATI2 deletion mutants tested. The top bar shows the potential binding sites predicted by the ANCHOR2 software in blue [50]. The designation of the different parts of the protein is consistent with evidence in [30]. IDR, intrinsically disordered region; TM, transmembrane segment; C-term, C-terminal region. (B) Yeast two-hybrid interactions between the ATI2 versions shown in (A) and NcGN^AGO1^. The interaction tests were done as described in Figure 3A.

**Figure 5.**
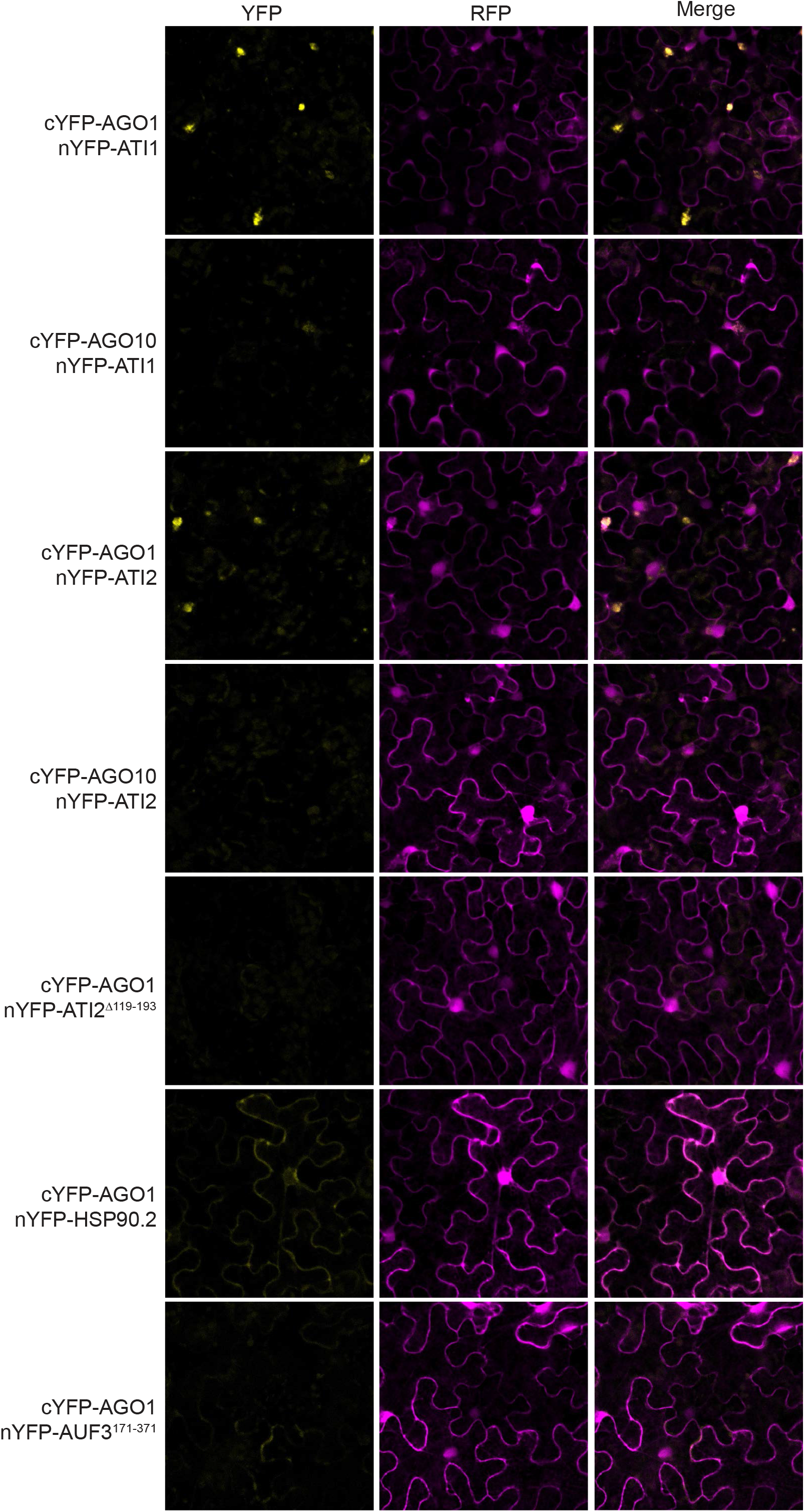
Bimolecular fluorescence complementation assays of protein-protein interactions involving full length AGO1. Confocal images bimolecular fluorescence complementation (BiFC) assays to probe the AGO1-ATI1 interaction *in vivo* upon transient expression in *Nicotiana benthamiana*. The C-terminal half of YFP was fused to the N-terminus of AGO1 or AGO10. The N-terminal half of YFP was fused to ATI1, ATI2, Hsp90.2 or the receptor part of AUF3 (AUF3^171-371^), excluding the F-box and small N-terminal extension of the protein. Both fused halves of YFP were expressed from the same plasmid that also expresses free RFP. Interaction is detected by fluorescence intensities from reconstituted YFP exceeding those obtained with the ATI2^Δ119-193^ negative control. RFP fluorescence serves as a control for transformation. Merged channel images demonstrate the presence of yellow fluorescence only in transformed cells. Scale bar, 50 μm.

### The N-coil is of special importance for interactions via the NcGN

We next focused our attention on the possible importance of the 14-aa N-coil for interaction, among other reasons because two mutations with functional impact have been identified in the N-coil of AGO1 (G186R, G189E) [38, 39]. We initially screened the interactors for the ability to interact with the NcGN carrying the G186R mutation. This point mutation in the N-coil resulted in strongly reduced interaction with most candidates as measured by β-galactosidase activity originating from *lacZ* expressed under UAS_Gal_ control in the reporter strain (Figure 6A). Indeed, of five proteins tested (ATI1, ATI2, TCTP, AUF1 and Hsp90) only Hsp90 and AUF1 showed clear interaction upon deletion of the N-coil (Supplemental Figure S3). Although the N-coil was clearly required for interaction with several proteins, it was not sufficient to yield a positive two-hybrid interaction with any candidate other than Hsp90.3 (Figure S3). On the one hand, these results point to a special importance of the small, 14-aa N-coil in several protein-protein interactions. On the other hand, they also show that NcGN^AGO1^ engages in protein-protein interactions independently of the N-coil via the globular N domain. We therefore aimed to characterize these two modes of protein-protein interaction in further detail using the NcGN^AGO1^ interactions with ATI1/2 and AUF1 as model systems.

**Figure 6.**
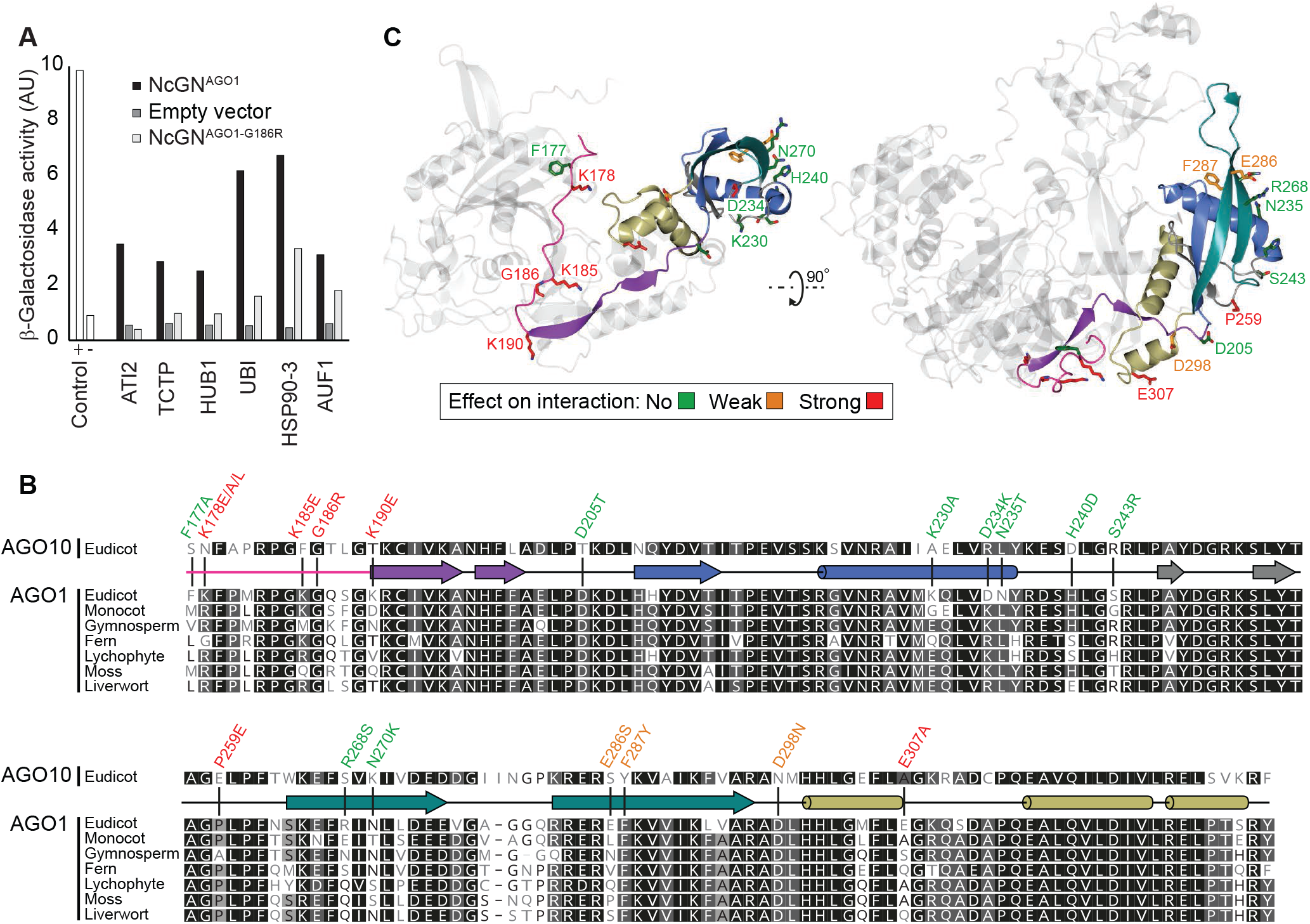
The N-coil is particularly important for specific interactions with NcGN^AGO1^. (A) β-galactosidase activity measured in an *ortho*-nitrophenyl-β-galactoside (ONPG) assay conducted with total protein lysates prepared from yeast strains expressing each candidate fused to Gal4-AD and AGO1-NcGN-Gal4-BD, AGO1-NcGN^G186R^-Gal4-BD, or unfused Gal4-BD (empty). y-axis shows β-galactosidase activity in arbitrary units. The yeast Swi2-Swi6 interaction was used as positive control, and co-expression of unfused Gal4-AD and Gal4-BD was used as negative control. (B) Sequence alignments of the arabidopsis paralogs AGO1 and AGO10, and of orthologs of arabidopsis AGO1 from plant species representing major phylogenetic groups of land plants. The species chosen from the different groups were as follows: Liverwort, *Marchantia polymorpha*; moss, *Ceratodon purpureus*; lycophyte, *Selaginella moellendorfii*; fern; *Ceratopteris richardii*; gymnosperm, *Pinus tabuliformis*; monocot, *Elaeis guineensis*; dicot, *Arabidopsis thaliana*. Residues selected for mutational analysis are marked on the alignment, and the effect of the indicated mutations on NcGN^AGO1^-ATI1/2 interaction is shown by color coding. (C) Alphafold ribbon model of arabidopsis AGO1 with the N-coil and N domain highlighted in rainbow colors. The residues studied by mutation are presented in sticks and colored according to the effect of mutation on ATI1/ATI2 interaction with AGO1-NcGN in the yeast two-hybrid assay. The underlying evidence from yeast-two hybrid spotting assays is shown in Supplemental Figure S4 and S5.

### The interactions of NcGN^AGO1^ with ATI1/ATI2 and with AUF1 require distinct sets of amino acids

To identify amino acid residues in NcGN^AGO1^ required for ATI1/2 interaction, we first inspected alignments between AGO1 orthologues from different plant species and between arabidopsis AGO1 and its closest paralogue, AGO10 (Figure 6B). This analysis revealed multiple sites of variation between AGO1 and AGO10 concentrated in the N-coil and in distinct sites in the GN-domain (Figure 6B). Several of these sites corresponded to surface-exposed patches, consistent with an involvement in protein-protein interaction (Figure 6C). We therefore constructed a series of point mutations in NcGN^AGO1^ residues variable between AGO1 and AGO10, and tested the interaction in the yeast two-hybrid system with ATI1 and ATI2. The results showed that multiple residues in the N-coil (K178, K185, K190) and in a few surface-exposed patches of the GN (P259, E307) are required for interaction with ATI1/2 (Figure 6C, Supplemental Figure S4-S5). In particular, mutation of either lysine residue to glutamic acid throughout the N-coil (K178E, K185E, K190E) disrupted the NcGN-ATI1/2 interaction.

We next used the same approach to identify residues in GN^AGO1^ important for the AUF1 interaction (Figure 7A,B). This effort revealed a distinct set of GN residues to be implicated in AUF1 interaction compared to ATI1/2 interaction. Specifically, none of the GN residues shown to be important for ATI1/2 interaction (P259, E307) had any impact on AUF1 interaction, while mutation of D205 and M304 strongly reduced AUF1 interaction, but had no effect on ATI1/2 interaction (Figure 7B). In addition, although mutation of the surface-exposed patch comprising residues K230, N234, T235 had no measurable effect on AUF1 interaction on its own, it enhanced the defect of the D205T mutant, indicating that it is involved in AUF1 interaction (Figure 7B, Supplemental Figure S4-S5). We conclude from these mutational analyses that NcGN^AGO1^ uses at least two distinct modes for binding to regulatory factors: one that depends critically on the N-coil as well as a subset (P259, E307) of GN residues, and one that is independent of the N-coil and requires a distinct subset (D205, M304) of GN residues.

**Figure 7.**
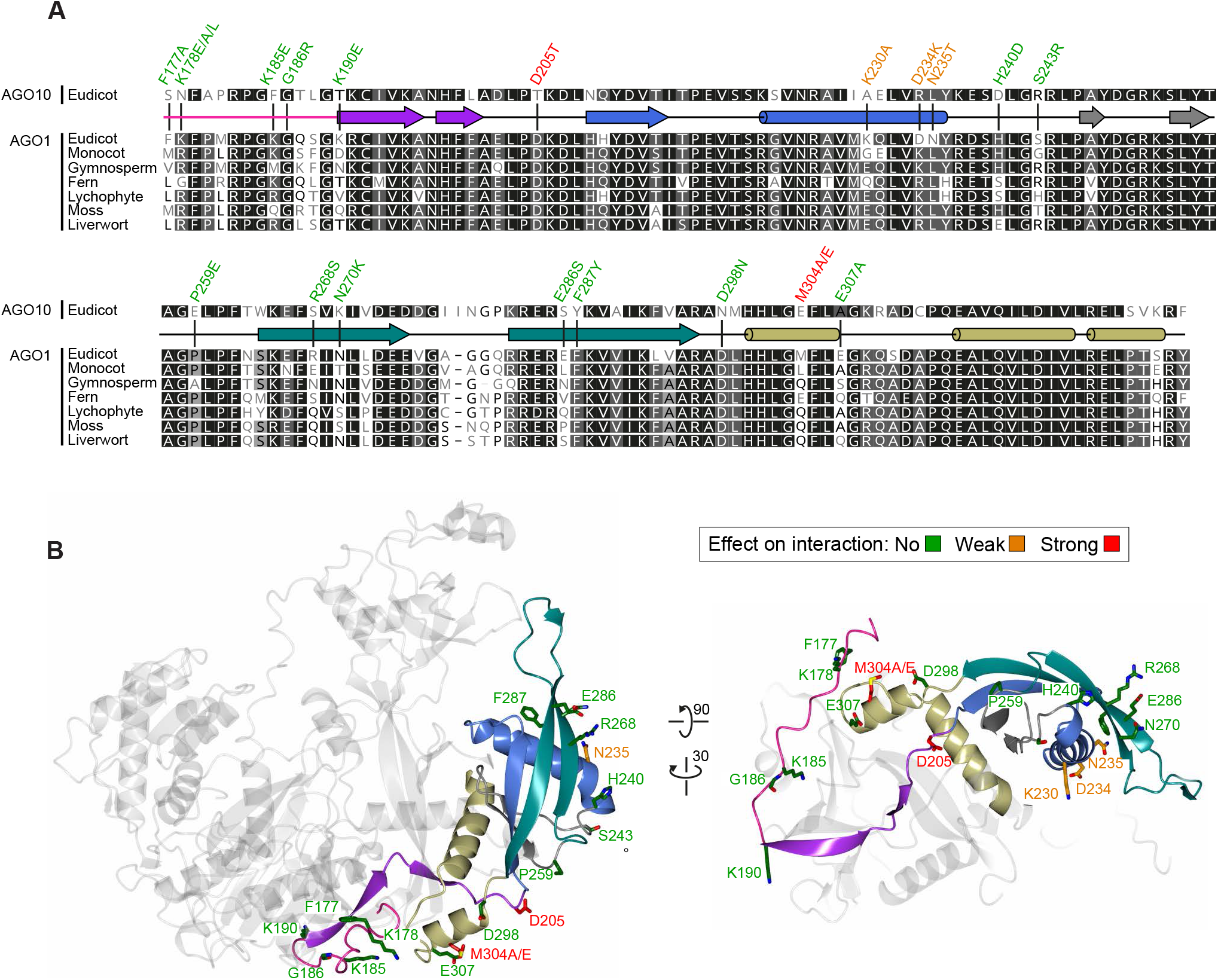
AUF1 interaction with NcGN^AGO1^ requires amino acid residues distinct from those involved in ATI1/2 interaction. (A) Alignments of arabidopsis AGO1 and AGO10, and of orthologs of arabidopsis AGO1, as described in Figure 6B. The effect of the indicated mutations on the NcGN^AGO1^-AUF1 interaction is shown by color coding. (B) Alphafold ribbon model of Arabidopsis AGO1 analogous to that shown in Figure 6C, but with views allowing visualization of amino acid residues in GN^AGO1^ necessary for NcGN^AGO1^-AUF1 interaction in yeast. The underlying evidence from yeast-two hybrid spotting assays is shown in Supplemental Figure S4 and S5.

### *The N-coil-dependent interaction mode and ATI1/2 influence the degradation rate of the NcGN^AGO1^* in planta

To probe the *in vivo* importance of the modes of NcGN^AGO1^ interaction identified by the combined mutational and interaction analysis in yeast, we took four properties revealed in this study into account. (1) Fusion of the NcGN^AGO1^ to YFP markedly increases the turnover rate of YFP. (2) Many NcGN^AGO1^ interactors mediate regulated proteolysis. (3) The majority of the identified interactors use the N-coil-dependent mode NcGN^AGO1^ interaction. (4) The only NcGN^AGO1^-specific interactor clearly shown to use the N-coil-independent mode of interaction is AUF1, an auxin-induced F-box protein whose expression is undetectable in the absence of auxin treatment [40]. We therefore set out to analyze the effect on the identified NcGN^AGO1^ interaction modes on expression of NcGN^AGO1^-YFP fusions with a particular emphasis on the N-coil-dependent mode. To this end, we first used CRISPR-Cas9 to generate a frameshift deletion allele of *ATI2* in the Col-0 ecotype. The resulting *ati2-3* mutant allele was combined with the *ati1-1* T-DNA allele to yield *ati1-1 ati2-3* double mutants. Western blot analysis with ATI2 [30] and ATI1 antibodies confirmed that both proteins were undetectable in *ati1-1 ati2-3* mutants, indicating that the *ati1-1 ati2-3* mutant is a double knockout (Figure 8A). We then generated transgenic lines expressing NcGN^AGO1^-YFP in both wild type and *ati1-1 ati2-3* mutants, and compared ratios between YFP protein and mRNA in lines that do not produce YFP siRNAs. These results showed modestly higher YFP protein/mRNA levels (~2-fold) in *ati1-1 ati2-3* compared to wild type (Figure 8B-D). Since the mRNA from these constructs is identical in both backgrounds, it is a fair assumption that their translation, i.e. synthesis rate, is unchanged. Thus, the most straightforward interpretation of these results is that ATI1/ATI2 are implicated in degradation of AGO1 via interaction with NcGN^AGO1^ *in vivo*. This is consistent with the involvement of ATI1/ATI2 in AGO1 degradation elicited by the poleroviral silencing suppressor P0 [31]. We note, however, that we cannot formally exclude an indirect effect of loss of ATI1/ATI2 on the translation of the NcGN^AGO1^-YFP mRNA.

**Figure 8.**
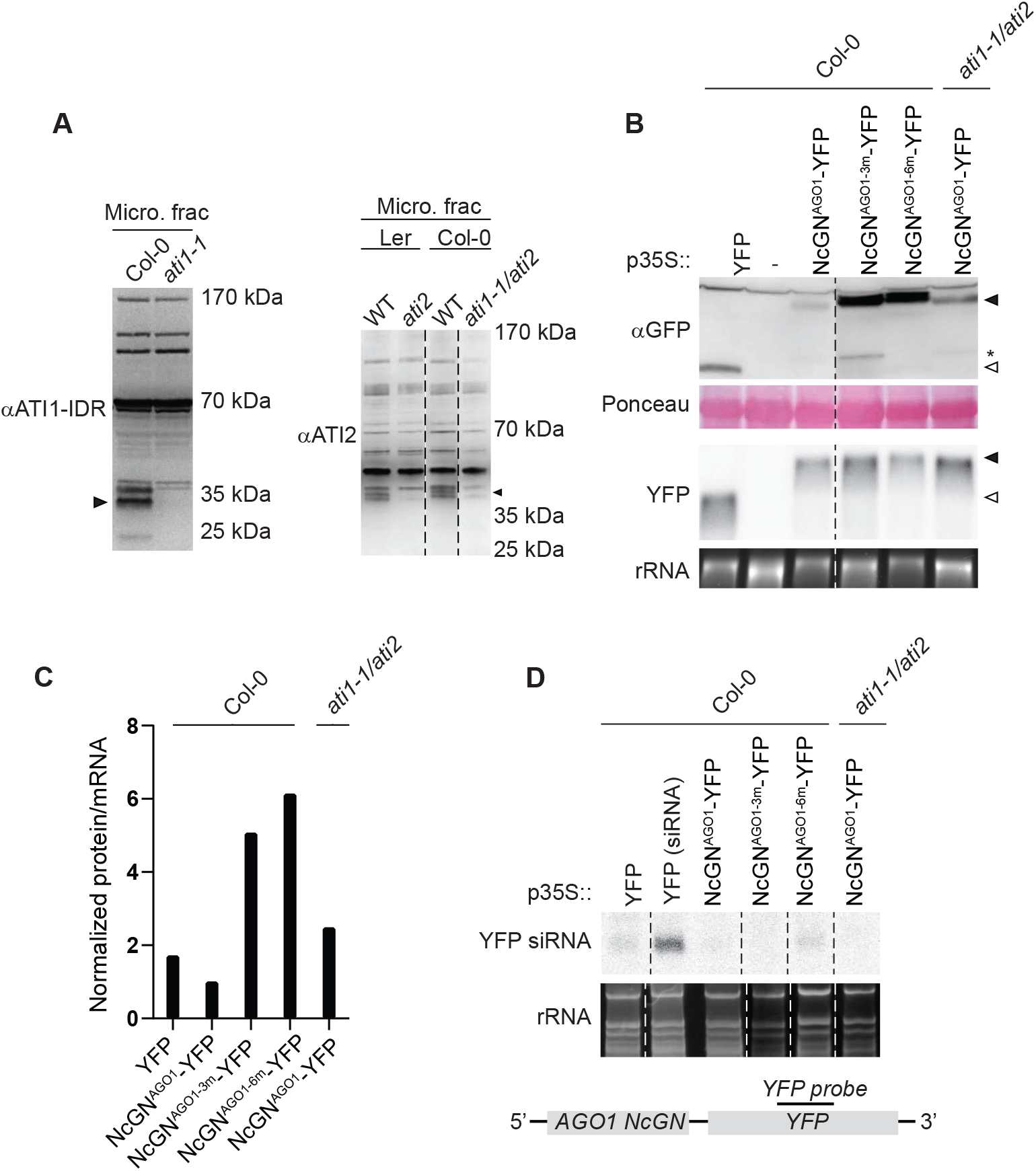
The NcGN^AGO1^ protein/mRNA ratio increases upon mutation of ATI1/2-interaction residues or of *ATI1/2*. (A) Microsome fractionation of *ati1-1* (left) and *ati2* (right) knock-out lines developed with antibodies recognizing the native proteins. Only membrane fractions were analyzed. Black arrowheads indicate the migration of the proteins in WT lines. (B) Levels of NcGN^AGO1^-YFP fusion proteins (black arrowhead) and YFP protein alone (white arrowhead) in Col-0 WT and *ati1-1 ati2-3* knock-out lines are shown in the upper panel. Corresponding mRNA levels are shown in the lower panel detected by a probe hybridizing to YFP as shown in (C). * indicates an assumed NcGN^AGO1^-YFP truncated protein. 3m, K178E/K185E/K190E; 6m, KKK/P259E/M304E/E307A. (C) Quantification of (A). Ratios are normalized to NcGN^AGO1(WT)^ expressed in the Col-0 WT background. (D) YFP siRNA levels of the NcGN^AGO1^ lines used in (B) and (C). siRNAs were detected by the YFP probe to document that differences in protein/mRNA ratios cannot be explained by different YFP siRNA levels in the transgenic lines. An NcGN^AGO1^-YFP line producing detectable levels of siRNAs was included as a positive control. The approximate position of the YFP probe is shown in the schematic representation of the *NcGN^AGO1^-YFP* fusion construct.

To test the importance of the identified NcGN^AGO1^ interaction modes more generally and without relying on mutation of specific interactors such as ATI1 and ATI2, we next engineered two point mutants in NcGN^AGO1^-YFP and expressed them in the Col-0 wild type background. The NcGN^AGO1-3m^-YFP mutant had mutations in all three lysine residues in the N-coil shown to be important for ATI1/ATI2 interaction (K178E/K185E/K190E). A second mutant, NcGN^AGO1-6m^-YFP, contained the same N-coil mutations as NcGN^AGO1-3m^-YFP, but in addition, it harbored two mutations in residues in the GN required for ATI1/ATI2 interaction (P259E, E307A), and one mutation in the residue with the strongest effect on AUF1 interaction (M304E). Both NcGN^AGO1-3m^-YFP and NcGN^AGO1-6m^-YFP mutants showed markedly higher protein/mRNA ratios than NcGN^AGO1-WT^-YFP when expressed in plants (Figure 8B,C). We draw two conclusions from the results of these reporter expression assays. First, in combination, the N-coil-dependent and -independent modes of protein-protein interaction contribute clearly to NcGN^AGO1^-YFP degradation *in vivo*. This conclusion supports the idea that these modes of interaction have biological relevance in the context of the full length AGO1 protein. Second, the N-coil-dependent mode alone also has a clear effect on NcGN^AGO1^-YFP degradation. The results do not, however, allow conclusions to be drawn on the possible implication of the N-coil-independent mode in NcGN^AGO1^-degradation, because compared to the N-coil-specific 3m-mutant, the 6m-mutant contained further mutations affecting both the N-coil-dependent and -independent modes.

## DISCUSSION

### The N-terminal part of AGO proteins as a hub for regulatory interactions

The protein-protein interaction assays conducted here strongly suggest that the N-terminal part of AGO1, including the N-coil and the globular N domain, provides important interaction sites for the regulation of AGO1 abundance, and perhaps other properties such as localization. A regulatory, rather than core biochemical, function of the globular N domain is consistent with the absence in this domain of mutations that abrogate miRNA and siRNA function of plant AGO proteins [1].

### Use of distinct modes of interaction for regulation of loaded and unloaded AGO1?

It is a question of considerable interest, not answered in the present study, what the biochemical and biological relevance may be of the two distinct interaction modes of NcGN^AGO1^ described here. The accompanying study shows that the N-coil is only accessible for interaction in the free (i.e. not bound to small RNA) state of AGO1, also referred to as the unloaded state [41]. Thus, the N-coil-dependent interaction mode is specific for the unloaded AGO form that is well known to be turned over rapidly *in vivo* in many organisms [42–45]. Together with the identification of N-coil-dependent interactors in the present study, this now opens interesting lines of research on understanding molecular mechanisms that direct AGO1 in the free state either towards RISC maturation (“loading”) or degradation. In this context, it is also worth noting that our identification of Hsp90 as an interactor of the N-coil suggests a simple explanation for why chaperone interaction protects unloaded AGO proteins from rapid proteolysis [42]. If the N-coil, and potentially other structural determinants with similar preferential exposure in the unloaded state of AGO, are concealed via chaperone interaction, chaperone binding to an AGO protein would preclude its interaction with the factors targeting its unloaded form for proteolysis. We also note that our identification of factors implicated in regulated proteolysis as interactors of the N-coil may provide a molecular explanation to the observation that the Arabidopsis AGO1 G186R mutant protein encoded by the *ago1-38* mutant allele is less abundant in membrane fractions than the wild type protein. The G186 residue is universally conserved in eukaryotic AGO proteins as any other residue in this position would lead to a steric clash with the Piwi domain in the small RNA-bound conformation [1, 46]. Thus, the bulky Arg side chain may partially dislocate the N-coil from the Piwi domain in the loaded conformation, making it accessible to interactors, including those concentrated in membrane compartments such as ATI1/2 [29, 30], ultimately leading to increased degradation rates.

With the identification of the N-coil-dependent interaction mode as exclusive to unloaded AGO1 [41], it becomes tempting to speculate whether the N-coil-independent mode is preferentially, or perhaps exclusively, used by small RNA-bound AGO1. Although this possibility remains unsupported by evidence at present, at least one consideration makes it an attractive hypothesis for further study. The N-coil-independent interactor identified here is AUF1, an F-box protein likely to form a functional Cullin1-based SCF ubiquitin ligase complex because of its verified interaction with the SCF core component ASK1 [47]. AUF1 is strongly induced by the plant growth regulator auxin [40], and many highly abundant miRNAs that exert their activity in AGO1-based RISCs target auxin receptors and auxin response transcription factors [48]. Thus, it is possible that an efficient auxin response requires rapid remodeling of the active miRNA population via exclusive degradation of the loaded AGO1-small RNA complexes. Such a mechanism would be reminiscent of the enhanced ubiquitin-proteasome-dependent turnover of human Ago2 that occurs as an element of genetic reprogramming driving T cell activation [49].

## AUTHOR CONTRIBUTIONS

SK conducted two-hybrid screening and initial mutational analysis of NcGN^AGO1^, further developed by AM. SB conducted pulse-labeling and BiFC assays. IMZS developed the ATI1 antibody, identified regions of ATI2 necessary for interaction with NcGN^AGO1^ in the two-hybrid system, and, together with SB, constructed and analyzed transgenic NcGN^AGO1^-YFP (and mutants thereof) in wild type and *ati1/ati2* double mutant backgrounds. EDO set up CRISPR-Cas9 engineering, isolated *ati2* mutant alleles in Col-0 and constructed *ati1/ati2* double mutants. CP conducted structural analyses.

## ACKNOWLEDGEMENTS

Theo Bølsterli and his team are thanked for plant care. Geneviève Thon is thanked for providing the PJ69-4A strain. This work was supported by a Hallas Møller Stipend (Hallas Møller 2010) and a project grant (NNF17OC0029194) from the Novo Nordisk Foundation, and Starting (ERC-2011-StG 282460 MICROMECCA) and Consolidator Grants (ERC-2016-CoG 726417 PATHORISC) from the European Research Council to PB.

## SUPPLEMENTAL FIGURE LEGENDS

**Supplemental Figure S1. Confirmed yeast-two hybrid interactions with NcGN^AGO1^.**

Candidate interactors of NcGN^AGO1^ found in the original screen fused to the activation domain (AD) of the yeast transcription factor Gal4 co-expressed with NcGN^AGO1^ fused to the binding domain of Gal4 (BD). The interaction between yeast Swi2 and Swi6 was used as positive control and empty vectors as negative control. 10-fold serial dilutions of yeast cells were spotted on both permissive (left) and selective (right) media.

**Supplemental Figure S2. Yeast two-hybrid analysis of interaction between ATI2, AUF1, AUF3, HUB1, HSP90.3, PP2C or UBI-like and NcGN^AGO4^ and NcGN^AGO10^.**

Candidate proteins fused to the activation domain (AD) of the yeast transcription factor Gal4 together were co-expressed with the NcGN domain of AGO4 or AGO10 fused to the binding domain of Gal4 (BD). The interaction between yeast Swi2 and Swi6 was used as positive control. 10-fold serial dilutions of yeast cells spotted on both both media selecting for presence of Gal4-AD/Gal4-BD plasmids (left), and selecting for interaction (right) are shown.

**Supplemental Figure S3. Yeast two-hybrid analysis of interaction between NcGN^AGO1^, GN^AGO1^ or extended N-coil and ATI1, ATI2, TCTP, AUF1 or HSP90.3**.

Candidate proteins fused to the activation domain (AD) of the yeast transcription factor Gal4 were co-expressed with NcGN^AGO1^, N-coil plus connecting β-strands (β-Nc), GN^AGO1^ fused to the DNA binding domain of Gal4 (BD). Co-expression of Gal4-AD fusions with unfused Gal4-BD (Empty) was used as a negative control. 10-fold serial dilutions of yeast cells spotted on both media selecting for presence of Gal4-AD/Gal4-BD plasmids (left), and selecting for interaction (right) are shown.

**Supplemental Figure S4. Identification of residues in NcGN^AGO1^ required for ATI1/2 and AUF1 interaction.**

Yeast-two hybrid analysis of ATI1 and ATI2 fused to the activation domain (AD) of the yeast transcription factor Gal4 co-expressed with the point mutants and truncated versions of NcGN^AGO1^ fused to the binding domain of Gal4 (BD). Yeast Swi2 and Swi6 interaction was used as positive control and empty vectors used as negative control. 10-fold serial dilutions of yeast cells were spotted on both permissive (left) and selective (right) media.

**Supplemental Figure S5.** Immunoblot analysis with α-Gal4-BD antibody of yeast strains expressing point mutants and truncated versions of AGO1-NcGN fused to the binding domain of GAL4 (BD). Empty vectors were used as negative control. Ponceau staining shows total protein.

## METHODS

### Plant material and growth conditions

The *Arabidopsis thaliana* Col-0 and Ler accessions were used in this study. All transgenic lines were constructed in accession Col-0. The *ati1-1* (SAIL404-D10, accession Col-0) and *ati2-2* (GT_5_4264 JIC SM line, accession Ler) insertion lines were obtained from the Nottingham Arabidopsis Stock Centre. Seeds were sterilized as described in [52] prior to being sown on Murashige-Skoog (MS) agar medium (4.3 g/L MS salts, 0.8 % agar, 1% sucrose). The seedlings for experiments were grown at constant temperature (21°C) and a 16-h light (80 μE m^−2^ s^−1^)/8-h darkness cycle. Transgenic lines and mutants were propagated in soil under greenhouse conditions.

### Yeast two-hybrid bait and libraries

A yeast two-hybrid bait plasmid was constructed by cloning an AGO1 cDNA fragment encoding residues Q165-Y335 (NcGN^AGO1^) into the pGBKT7 vector (Clontech). Three libraries were used for the yeast two-hybrid screen of candidates interacting with AGO1 NcGN: CD4-22, yeast two-hybrid cDNA library generated with oligo-dT primed mRNA (0.6-2.5 kb fragments) isolated from 3-day-old etiolated arabidopsis seedlings [51], distributed by Arabidopsis Biological Resource Center, ABRC); CD4-10, yeast two-hybrid library (>300 bp fragments) of random primed mRNA extracted from leaves and roots from mature arabidopsis plants, distributed by ABRC; MM, Matchmaker yeast two-hybrid library of arabidopsis cDNA purchased from Clontech. CD4-22 and CD4-10 libraries were converted from λ cDNA into plasmid libraries by transformation into BNN132 bacterial cells from which the plasmids were extracted. The yeast strain PJ69-4A[53] (*trp1-901, leu2-3, 112, ura3-52, his3-200, gal4Δ, gal80Δ, LYS2::GAL1-HIS3, GAL2-ADE2, met2::GAL7-lacZ*) was used for all two-hybrid screens and interaction tests.

### Yeast two-hybrid assays

Library transformation was performed as described by Clontech (Yeast Protocols Handbook) except that a 1h recovery step at 30°C was inserted between heat shock at 42°C and plating on selective media (synthetic defined (SD) medium) minus Trp, Leu, His (Clontech), supplemented with 1.5 mM 3-amino-1,2,4-triazol (3-AT, Sigma). Selection plates were left for 8 days at 30°C, and growing colonies were restreaked three times on selective media prior to plasmid isolation and transformation of *E. coli*. For plasmid corresponding to each yeast colony, 12 *E. coli* colony PCRs were run. All plasmids with different insert sizes were sequenced, and co-transformed with the NcGN^AGO1^ bait plasmid back into PJ69-4A to establish the set of candidate interactors shown in Table S1.

ATI1 was not recovered in any screen, but was tested in a directed two-hybrid assay in which cDNAs encoding ATI1 and ATI2 were cloned into the prey vector pGADT7 (Clontech) and co-transformed into PJ69-4A with the NcGN^AGO1^ bait plasmid. Additional bait vectors to test interactions with the NcGNs of AGO4 and AGO10 and with distinct parts of the AGO1 NcGN were also constructed in pGBKT7 [NcGN^AGO1^ (Q165-Y335), βNc^AGO1^ (S174-N197), GN^AGO1^ (P204-Y335), NcGN^AGO4^ (R41-L238) and NcGN^AGO10^ (V114-P275)]. Point mutations in NcGN^AGO1^ and GN^AGO1^ were introduced by site directed mutagenesis using Quickchange (Stratagene). Deletion variants of ATI2 were amplified from ATI2 (AT4G00355.1) CDS by overlap extension PCR using Phusion High Fidelity DNA polymerase (NEB) and subcloned into pCR™4/TOPO™ vectors (Invitrogen). Inserts were excised by appropriate restriction enzymes and ligated into the yeast prey plasmid pGADT7. The sequence of all final plasmids was verified by Sanger sequencing. All primers used are specified in Supplemental Table 2.

For co-transformations, a yeast culture of OD_600_=0.1-0.3 was freshly prepared for transformation by lithium acetate treatment by being resuspended sequentially in sterile water and a solution of TE/LiAc (11 mM tris-HCl pH 7.5, 1.1 mM EDTA pH 8, 110 mM LiAc). Prepared competent cells were mixed with 200 ng of prey DNA, 100 ng of bait DNA, and 50 μg of denatured Yeastmaker™ Carrier DNA (Clontech). The cells were incubated with PEG4000 buffer (40% [w/v] PEG4000, 100mM LiAc, 10mM Tris-Hcl, 1mM EDTA) for 30 min at 30°C before DMSO was added. The cells were then heat shocked for 15 min at 42 °C. After incubation, the cells were pelleted and resuspended in 100 μl 0.9 % NaCl and grown on SD minimal medium supplemented with drop-out medium lacking tryptophan and leucine for 2-3 days at 30°C. Cultures of colonies autotrophic for tryptophan and leucine were normalized according to their OD_600_ and spotted as a 10X serial dilution on SD minimal medium plates lacking Trp/Leu and Trp/Leu/His including positive and negative controls on all plates. 1.5 mM 3-AT was added to the Trp/Leu/His plates to inhibit leaky expression of *HIS3*. Cells were grown for 6-7 days at 30°C.

### Antibodies

Antibodies specific for the DNA-binding (BD) and activation domains of Gal4 (TA) were purchased from Santa Cruz Biotechnologies (anti-Gal4-DBD (sc-510) and anti-Gal4-TA-C10 (sc-429)). Polyclonal rabbit ATI2 antibodies were raised previously and were used as described [30]. To raise ATI1 antibodies, the N-terminal IDR of ATI1 comprising residues 1-180 was expressed as a GST-fusion (GST-ATI1^1-180^) in *E. coli* and purified as described [30]. 0.3 mg of protein was used per rabbit for immunization of 2 rabbits (Eurogentec). Serum collected from one rabbit was affinity purified against GST-ATI1^1-180^. GFP-Trap beads from ChromoTek were used to IP YFP fusions and an anti-GFP polyclonal antibody from AbCam (#ab290) was used for detection of YFP and YFP fusions.

### Protein extraction and analysis from yeast

Protein for western blotting was extracted from 10 ml overnight cultures grown in selective media. Pelleted cells were dissolved in 20 % (v/v) trichloroacetic acid (TCA) and lysed with glass beads by vortexing for 40 seconds on a Fastprep-24 5G (MP Biomedicals). Glass beads were removed and protein was pelleted by centrifugation. The protein pellet was washed in 80 % acetone and boiled in Laemmli buffer before being subjected to SDS-PAGE. Protein was transferred to a nitrocellulose membrane and incubated o/n with anti-Gal4 AD (1:1000 monoclonal mouse anti-Gal4 TA in 5 % milk in 1xPBS-T [0.05 % Tween-20], Santa Cruz Biotechnology) or anti-Gal4 BD antibody (1:2000 monoclonal mouse anti-Gal4 DBD in 5 % milk in 1xPBS-T [0.05 % Tween-20], Santa Cruz Biotechnology). Antimouse IgG conjugated to horseradish peroxidase was used as secondary antibody and the blots were developed with homemade enhanced chemiluminescence (ECL) (2.5 mM luminol, 0.4 mM p-coumaric acid, 0.1 M Tris–HCl (pH 8.5), 0.018 % H_2_O_2_), 1 tablet NBT-BCIP in water (Roche), or SuperSignal™ West Femto Maximum Sensitivity Substrate (Thermo Fisher) using films or digital images taken with a Sony A7S camera.

### *Construction of* ati2 *knockout mutants using CRISPR-Cas9*

pKI1.1R (Addgene plasmid #85808; http://n2t.net/addgene:85808; RRID:Addgene_85808) [54] was used to introduce sgRNAs (see Supplemental Table 2) targeting the *ATI2* gene in order to generate mutations, in an *ati1-1* background. Two plasmids were generated, as described in [54], by ligation into pKI1.1R: pKI1.1R-ati2-A and pKI1.1R-ati2-C. Plasmid A contains a sgRNA targeting the sequence AAGATGAAGCAGCGACGCG in ATI2, and plasmid B contains a sgRNA targeting the sequence CCTGGTCCTAAACCAGTAG located 100 bp downstream of the target site for plasmid A. The exact sequences of the oligos used for ligation into the vector are listed in Supplemental Table 2.

The *ati2* CRISPR knockout line was generated by transformation of *ati1-1* plants with a mix of *Agrobacterium tumefaciens* strains bearing the pKI1.1-ati2-A and the pKI1.1R-ati2-C constructs. The two plasmids were transformed together into plants to induce larger deletions detectable by PCR via gel shift, as the two guides were placed with ~100 bp distance. Selection of transformants was done by germinating T1 seeds on MS-agar supplemented with 25 μg/ml hygromycin. Transformants were transferred to soil and plants carrying deletions were identified by PCR (primers in Supplemental Table 2). PCR products were visualized in a 2% agarose gel, and plants showing a gel shift were selected. PCR products from these plants were sequenced by Sanger sequencing to isolate plants in which the deletion caused a frameshift mutation. In T2, plants homozygous for the deletion were detected by PCR and gel shift followed by Sanger sequencing to confirm homozygosity. In T3, knockout of *ATI2* was confirmed by Western blot of microsome fractions from selected lines probed with an ATI2-specific antibody. Furthermore, to select the plants no longer expressing Cas9 in T3, plants homozygous for the deletion in ATI2 were genotyped for the presence of the plasmid. The *ati2* knockout allele used in this paper harbors an 89 bp-deletion causing a premature stop codon.

### Microsome fractionation of ati1-1/ati2 knockout lines

14-day-old seedlings were flash frozen in liquid nitrogen and ground to a fine powder. 0.2 g of powder was dissolved in 1.2 ml of microsome buffer (50 mM MOPS, 0.5 M sorbitol, 5 mM EGTA, complete protease inhibitor (Roche) at pH 7.6). Samples were spun at 8000 *g* for 10 min at 4°C. Supernatants were transferred to new tubes and repeatedly spun at 8000 *g* until no pellet was visible. Supernatants (‘total extracts’) were spun at 100□000 *g* for 30 min at 4°C in a Beckman Optima XP ultracentrifuge. Pellets were resuspended in microsome buffer (50 mM MOPS, 0.5 M sorbitol, 5 mM EGTA, complete protease inhibitor (Roche)) and repelleted by centrifugation at 100□000 *g* for 30 min at 4°C. Laemmli buffer was added to all fractions before boiling the samples and loading them onto a 4-12% Bis-Tris Criterion™ acrylamide gel (Bio-Rad) and run in MOPS buffer (50 mM MOPS, 50 mM Tris, 0.1% SDS, 1 mM EDTA pH 7.7). Proteins were blotted onto a nitrocellulose membrane.

To probe for ATI2, the membrane was blocked for 5 min with 2 % BSA, washed three times in 1x PBS-T (0.05% Tween-20) buffer and incubated o/n with anti-ATI2 antibody (1:500 in PBS-T with 2% BSA). The membrane was then washed three times in PBS-T buffer and incubated with HRP-conjugated anti-rabbit antibody (1:10,000, Sigma) for 4 h. After an additional three washes in PBS-T, blots were developed as described above. To probe for ATI1, blots were first blocked in 3% milk in PBS-T for 30 min, washed 3 times in PBS-T, then incubated with ATI1 antiserum at 1:250 dilution in 3% milk (PBS-T) overnight at 4°C, and finally developed as described for ATI2 probing.

### Construction of transgenic lines expressing NcGN^AGO1^-YFP versions

C-terminal YFP fusions of AGO1-NcGN were made using a genomic DNA fragment encoding AGO1-NcGN in the vector pCAMBIA230035SU [55] modified to encode a C-terminal YFP. Exonic fragments containing mutated residues of interest (K178E, K185E/K190E, P259E, M304E/E307A) were amplified from already constructed yeast two-hybrid plasmids pGBKT7 or plant expression plasmids pCAMBIA3300U containing the relevant mutations, and combined by USER cloning [55] to give the final *NcGN^AGO1-3m^*, (K178E/K185E/K190E), and *NcGN^AGO1-6m^* (K178E/K185E/K190E/P259E/M304E/E307A) versions fused to YFP in pCAMBIA230035SU. The USER cloning was done with KAPA HiFi Hotstart Uracil□+□Readymix (Roche) and USER compatible primers specified in Supplemental table 2. NcGN^AGO1^-YFP, AGO1 NcGN^AGO1(KKK)^-YFP and NcGN^AGO1(6m)^-YFP were transformed into Col-0. NcGN^AGO1^-YFP was also transformed into the *ati1-1/ati2-3* line. Primary transformants (T1) were selected on MS agar medium supplemented with 100 μg/ml ampicillin and 50 μg/ml kanamycin. Transformed plants were transferred to soil after approximately 10 days of growth on selective medium. For every construct, lines with a single T-DNA locus and comparable protein expression levels were selected in T2, and descendants homozygous for the insertion were identified in T3.

### Protein stability of NcGN^AGO1^

NcGN^AGO1^-YFP expressing plants were analyzed in the second generation (T2). 50 mg of ground tissue from 10-day old sterile-grown seedlings was dissolved in 150 μl Laemmli buffer and heated at 75°C for 5 min. Protein samples were loaded in a 4-20 % Criterion™ TGX^TM^ Precast Midi Protein Gel (Bio-Rad) and subsequently transferred onto a nitrocellulose membrane. NcGN^AGO1^-YFP and YFP were detected by anti-GFP antibody (Abcam, 1:5000 in 5 % milk in 1x PBS-T [0.05 % tween20]). Band intensities were quantified with Adobe Photoshop v23.5. For quantification of signal intensity in protein bands, background levels in the immediate vicinity of each band were subtracted from the band intensity due to unequal background levels across the membrane. For quantification of mRNA signals, the same background subtraction was applied to all bands. mRNA levels were further normalized to ribosomal RNA levels (rRNA) in each lane before each protein/mRNA ratio was calculated. The protein/mRNA ratio of the NcGN^AGO1^-YFP sample was arbitrarily set to 1, and the ratios of all other samples were calculated relative to this sample.

### RNA extraction and RNA blot analysis

Total RNA was extracted from 100 mg tissue of 10-day old seedlings in 1 ml of TRI Reagent (#T9424, Sigma Aldrich Denmark A/S) according to the manual provided by the manufacturer. The RNA was dissolved in 50% formamide for small RNA blots or sterilized water for further polysaccharide precipitation for RNA blots for mRNA detection. Polysaccharides were precipitated for 30 min on ice by addition of 1/10 volume cold 96 % ethanol and 1/30 volume 2.5 M NaOAc (pH 5.2). The RNA from the supernatant was subsequently precipitated at −20 °C for 20 min by the addition of 1/10 volume 2.5M NaOAc (pH 5.2) and 2.5 volumes 70 % ethanol. The pellet was washed in 70 % ethanol after which it was resuspended in sterile water. For small RNA blots, 10 μg of total RNA was loaded on an 18 % denaturing polyacrylamide gel, blotted onto a Hybond Nx membrane where it was chemically cross-linked with the soluble carbodiimide EDC [56]. siRNAs were detected with a YFP long probe (primers listed in Supplemental Table 2) labeled with ^32^P-dCTP prepared according to the manual of the Prime-a-Gene Labeling System (Promega). The blot was hybridized overnight at 42 °C with the probe before being washed with 2xSSC, 2 % SDS at 42 °C. For detection of YFP mRNAs, 6 μg of polysaccharide precipitated total RNA was loaded on a 1% agarose/formaldehyde gel. The RNA was transferred to a Hybond Nx membrane and UV cross-linked at 254nm. The ^32^P-dCTP labeled YFP long probe was used at 42°C for annealing to the mRNA o/n and washed with 2xSSC, 0.1 % SDS at 65 °C. The membranes were exposed to a phosphoscreen and imaged on a Typhoon FLA 7000 scanner (GE Healthcare).

### Pulse labeling

Seedlings were grown for 7 days on MS agar. For each time point, 8 seedlings were transferred to 1.5 ml liquid MS in a 12 well culture plate and grown for two more days. EasyTag™ Express35S Protein Labeling-Mix (Perkin Elmer) was then added to a final count of 50 μCi/ml and at given time points, seedlings were briefly washed in water and flash frozen in liquid nitrogen. The tissue was ground to a fine powder before it was solubilized in 500 μl IP buffer (50 mM Tris-HCl pH 7.5, 150mM NaCl, 10 % glycerol [v/v], 5mM MgCl_2_, 0.1% Nonidet P40 [v/v], 4mM DTT, 2 tablets complete EDTA-free protease inhibitor (Roche)) followed by 5 min incubation at 4°C. The lysate was cleared by centrifugation at 16,000 *g* for 5 min at 4 °C before it was immobilized on 20 μl GFP-Trap beads (ChromoTek) for 30 min at 4 °C. The beads were transferred to an empty gravity column and washed with 1.5 ml IP buffer. Immunoprecipitated protein was released from the beads in Laemmli buffer by heating at 75 °C for 5 min.

### Pulse quantification

After Western blot development the nitrocellulose membranes were exposed to a storage phosphor screen and the screen was imaged using a Typhoon FLA 7000 scanner (GE Healthcare). Total lysate, IP Western blot, and ^35^S signals were quantified using Image J and ^35^S signal/IP Western blot signal were plotted against pulse time. The pulse data was fitted to an equation of the type,

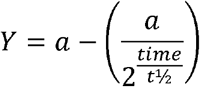

where *t*_1/2_ is the half-life of the immuno-precipitated protein and *a* is the maximum ^35^S value of radioactivity per protein that the sample will converge to when the pulse has reached equilibrium, i.e. when all unlabeled protein has been degraded and the newly synthesized proteins are radio-labelled. The data analysis was done using Non-linear Least Square (NLS) in R (version 4.2.0).

### Alignments and structural analyses

AGO1 sequences from diverse clades were aligned to arabidopsis AGO1 using Clustal Omega and presented with Boxshade coloring [57]. The following sequences given as “Clade | species | sequence” were aligned: Eudicot | *Arabidopsis thaliana* | O04379 (AGO1) and Q9XGW1 (AGO10), Monocot | *Elaeis guineensis* | A0A6J0PHR6, Gymnosperm | *Pinus tabuliformis* | AJP06229.1, Fern | *Ceratopteris richardii* | KAH7294104.1, Lycophyte | *Selaginella moellendorffii* | XP_002984845.2, Moss | *Ceratodon purpureus|* KAG0569476.1, Liverwort | *Marchantia polymorpha* | BAV72133.1. The predicted AlphaFold structure of arabidopsis AGO1 (AF-O04379-F1-model_v2) was used for highlighting the positions of amino acid residues chosen for mutational analysis. All the presented structural models were made using CCP4mg [58].

## ACKNOWLEDGEMENTS

We thank Theo Bölsterli and Rene Hvidberg and their teams for plant care.

