## Supplemental Figures for "The N-coil and the globular N-terminal domain of plant ARGONAUTE1 are interaction hubs for regulatory factors"

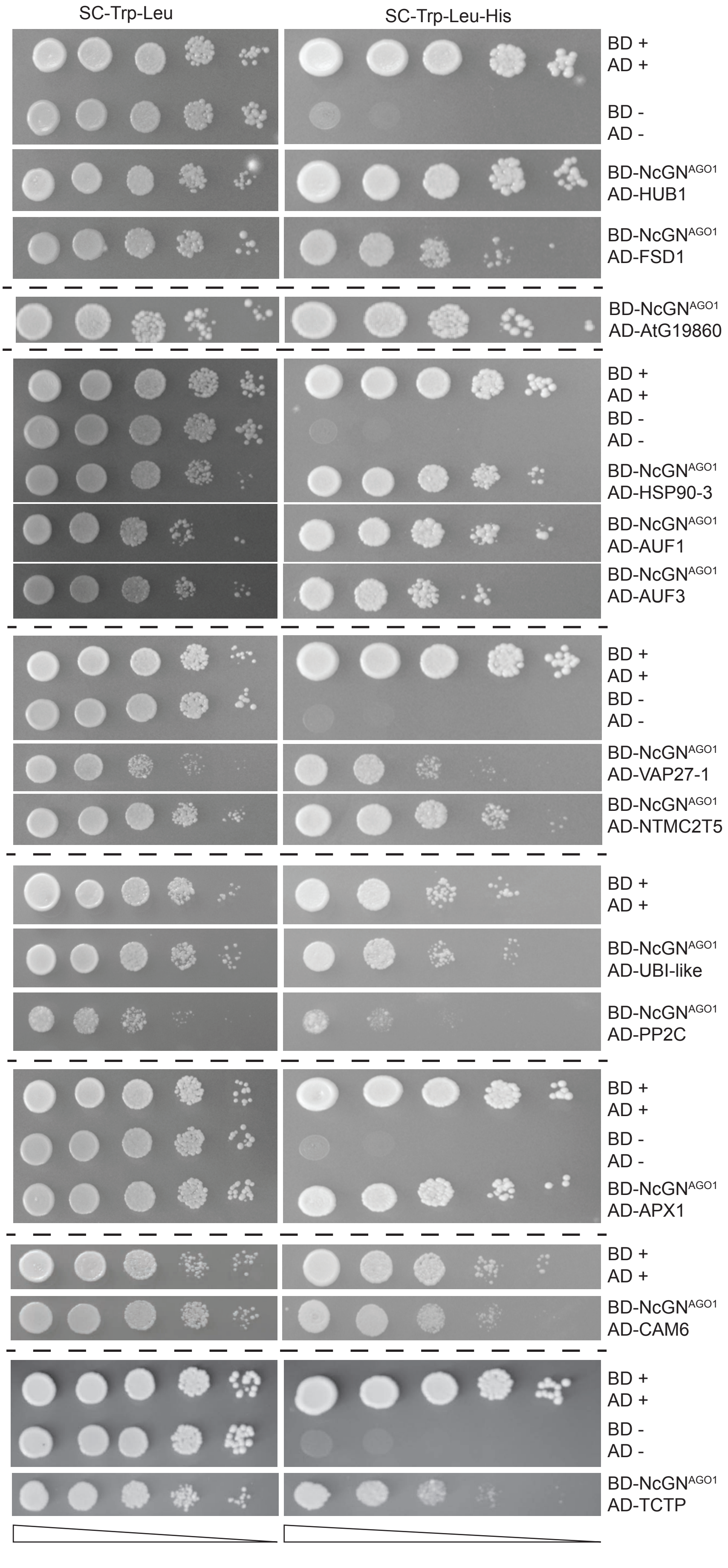


**Supplemental Figure S1. Confirmed yeast-two hybrid interactions with NcGN$^{AGO1}$.** Candidate interactors of NcGN$^{AGO1}$ found in the original screen fused to the activation domain (AD) of the yeast transcription factor Gal4 co-expressed with NcGN$^{AGO1}$ fused to the binding domain of Gal4 (BD). The interaction between yeast Swi2 and Swi6 was used as positive control and empty vectors as negative control. 10-fold serial dilutions of yeast cells were spotted on both permissive (left) and selective (right) media.

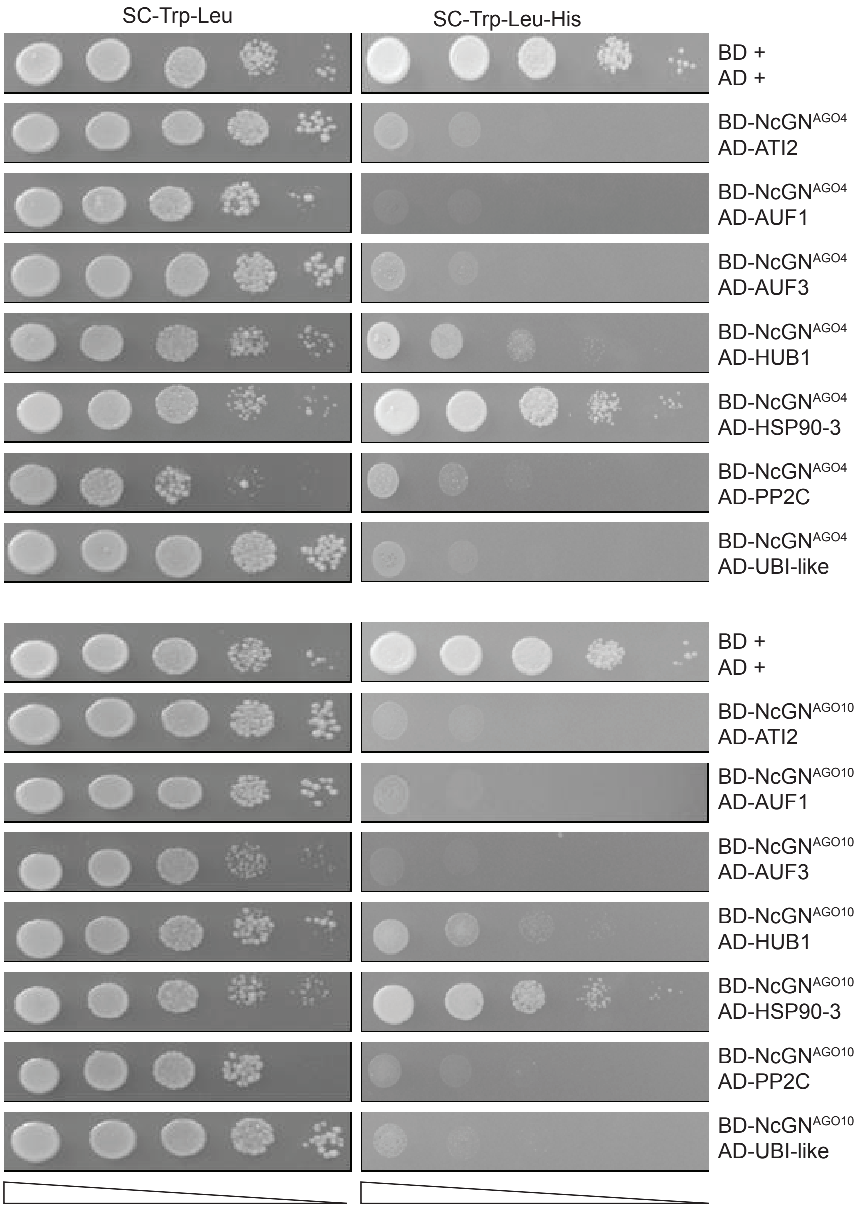


**Supplemental Figure S2. Yeast two-hybrid analysis of interaction between ATI2, AUF1, AUF3, HUB1, HSP90.3, PP2C or UBI-like and NcGN$^{AGO4}$ and NcGN$^{AGO10}$.**
Candidate proteins fused to the activation domain (AD) of the yeast transcription factor Gal4 together were co-expressed with the NcGN domain of AGO4 or AGO10 fused to the binding domain of Gal4 (BD). The interaction between yeast Swi2 and Swi6 was used as positive control. 10-fold serial dilutions of yeast cells spotted on both both media selecting for presence of Gal4-AD/Gal4-BD plasmids (left), and selecting for interaction (right) are shown.

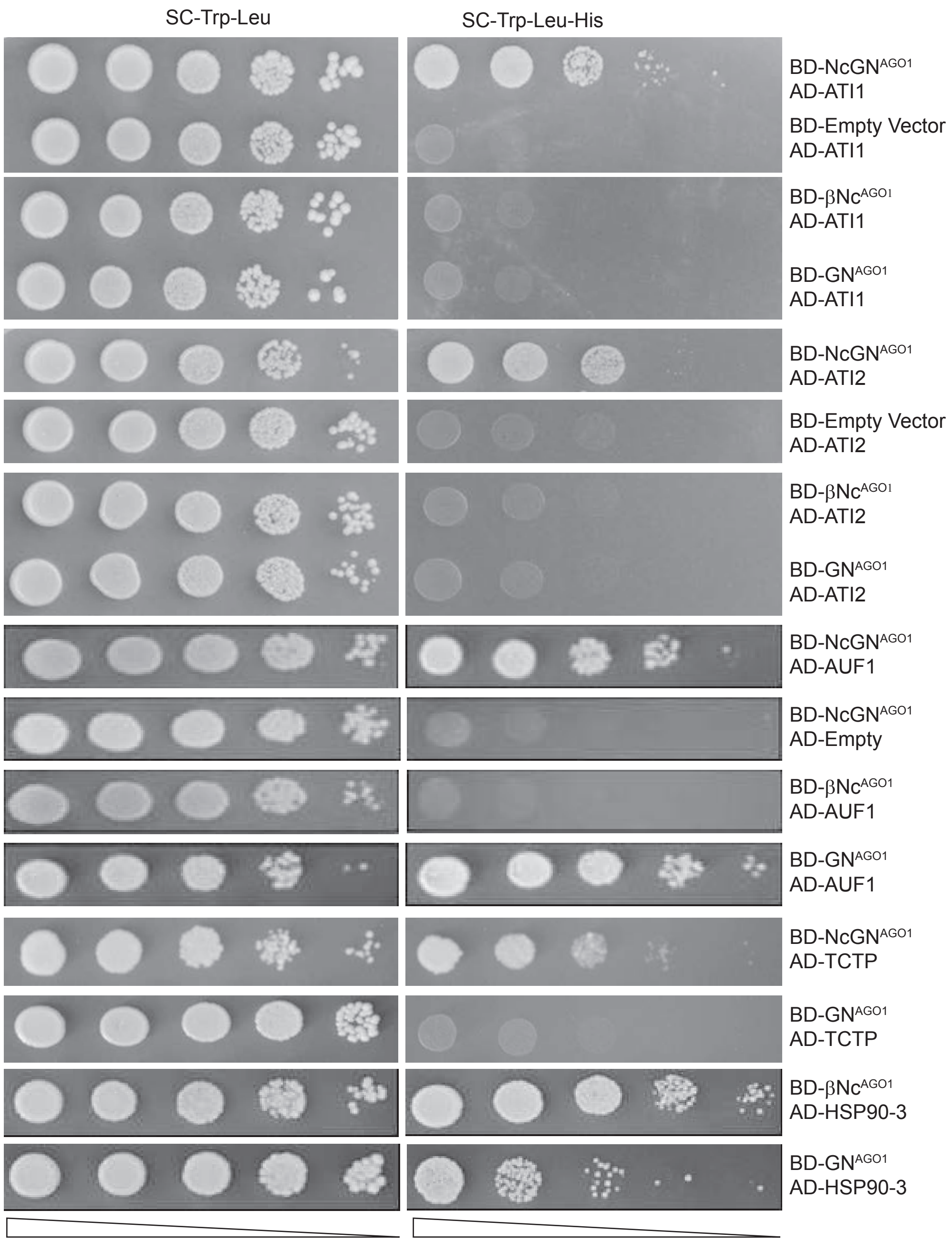


**Supplemental Figure S3. Yeast two-hybrid analysis of interaction between NcGN$^{AGO1}$, GN$^{AGO1}$ or extended N-coil and ATI1, ATI2, TCTP, AUF1 or HSP90.3**.

Candidate proteins fused to the activation domain (AD) of the yeast transcription factor Gal4 were co-expressed with NcGN$^{AGO1}$, N-coil plus connecting β-strands (β-Nc), GN$^{AGO1}$ fused to the DNA binding domain of Gal4 (BD). Co-expression of Gal4-AD fusions with unfused Gal4-BD (Empty) was used as a negative control. 10-fold serial dilutions of yeast cells spotted on both media selecting for presence of Gal4-AD/Gal4-BD plasmids (left), and selecting for interaction (right) are shown.

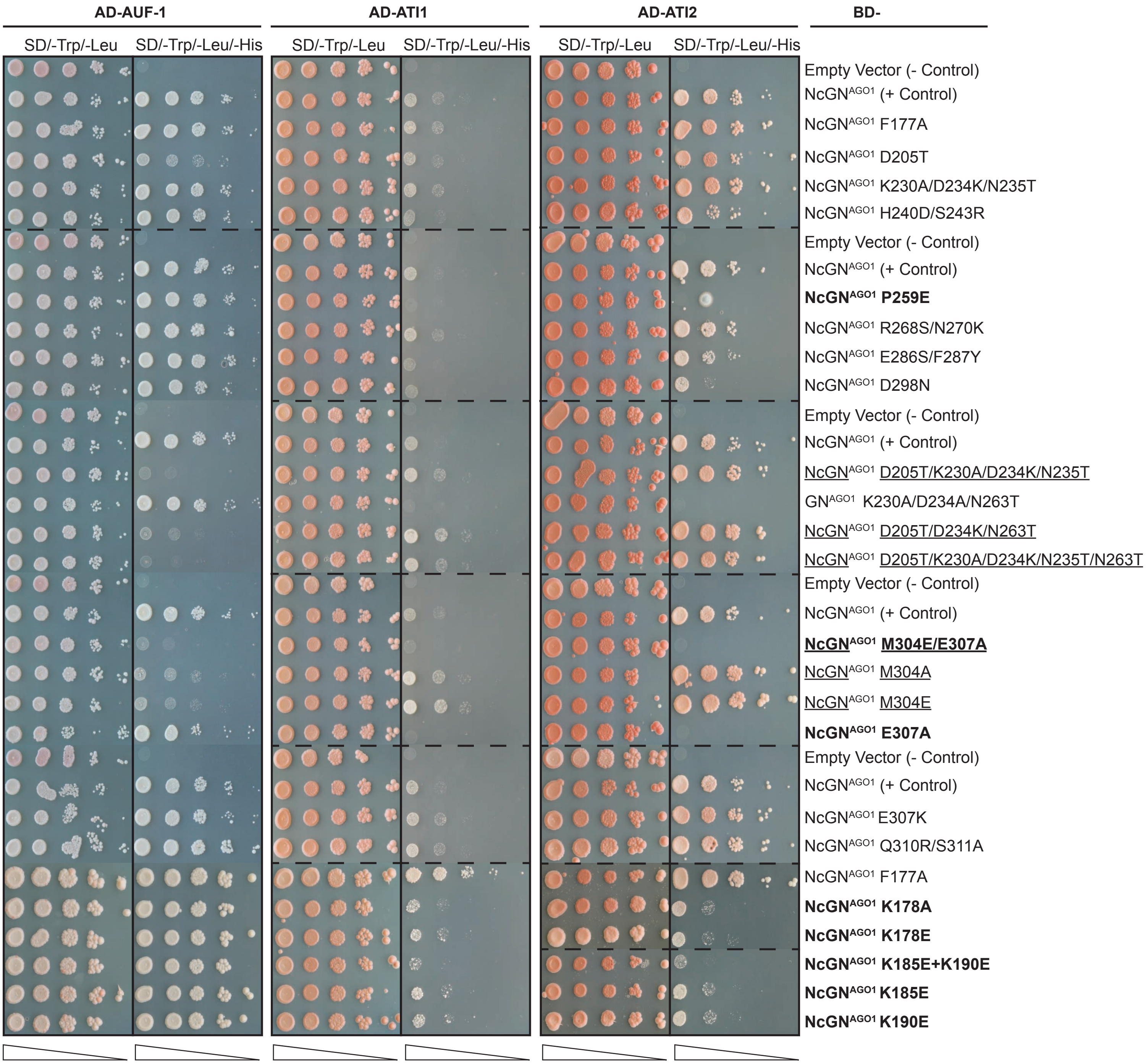


**Supplemental Figure S4. Identification of residues in NcGN$^{AGO1}$ required for ATI1/2 and AUF1 interaction.**
Yeast-two hybrid analysis of ATI1 and ATI2 fused to the activation domain (AD) of the yeast transcription factor Gal4 co-expressed with the point mutants and truncated versions of NcGN$^{AGO1}$ fused to the binding domain of Gal4 (BD). Yeast Swi2 and Swi6 interaction was used as positive control and empty vectors used as negative control. 10-fold serial dilutions of yeast cells were spotted on both permissive (left) and selective (right) media.

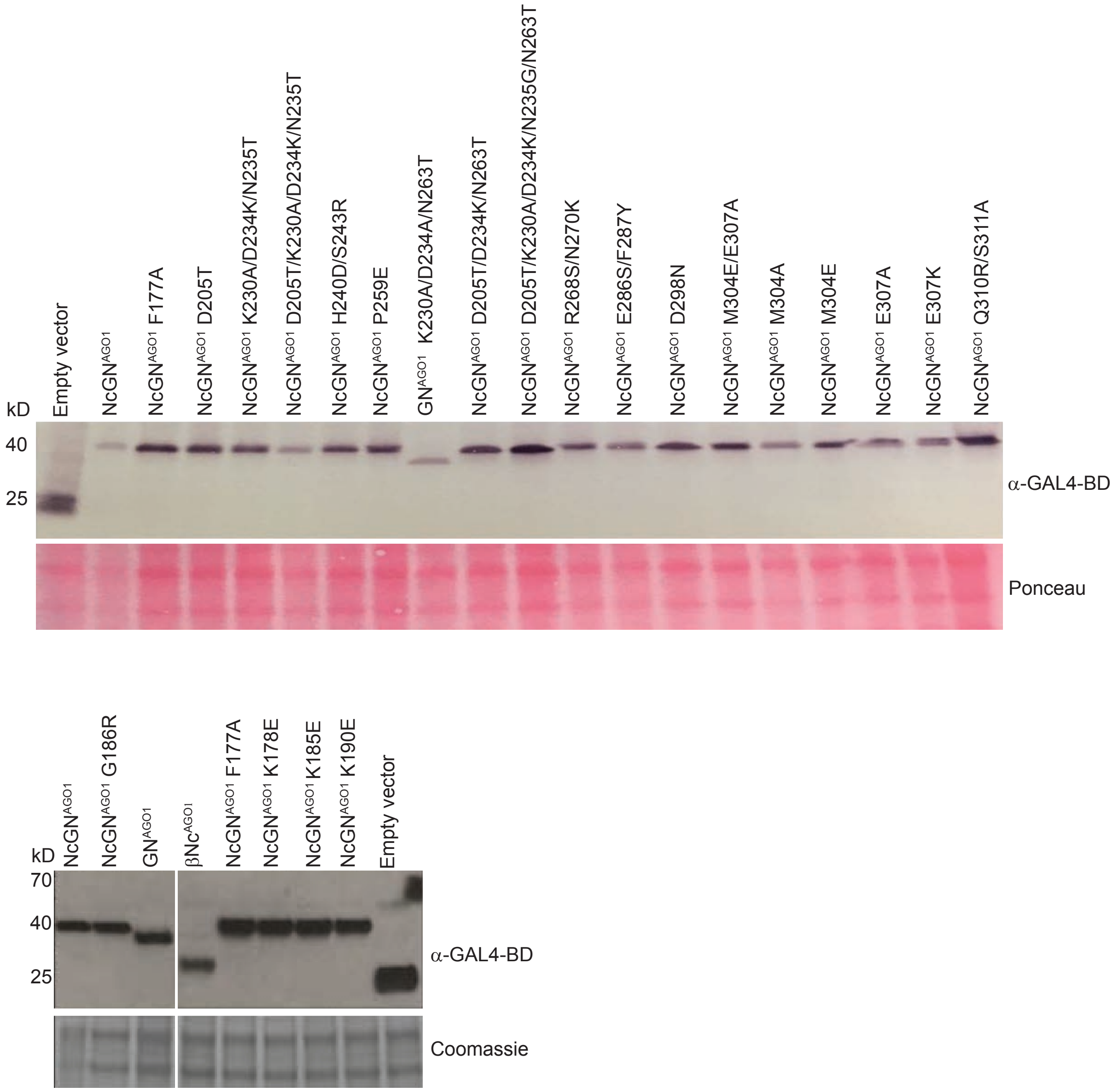


**Supplemental Figure S5.** Immunoblot analysis with α-Gal4-BD antibody of yeast strains expressing point mutants and truncated versions of AGO1-NcGN fused to the binding domain of GAL4 (BD). Empty vectors were used as negative control. Ponceau staining shows total protein.
